# Chidamide increases the sensitivity of non-small cell lung cancer to crizotinib by decreasing c-*MET* mRNA methylation

**DOI:** 10.1101/2020.03.28.012971

**Authors:** Nan Ding, Abin You, Wei Tian, Liankun Gu, Dajun Deng

## Abstract

**Introduction:** Crizotinib is a kinase inhibitor targeting c-MET/ALK/ROS1 used as the first-line chemical for the treatment of non-small cell lung cancer (NSCLC) with *ALK* mutations. Although c-MET is frequently overexpressed in 35-72% of NSCLC, most NSCLCs are primarily resistant to crizotinib treatment.

**Method:** A set of NSCLC cell lines were used to test the effect of chidamide on the crizotinib sensitivity *in vitro* and *in vivo*. Relationships between the synergistic effect of chidamide and c-MET expression and RNA methylation were systemically studied with a battery of molecular biological assays.

**Results:** We found for the first time that chidamide could increase the crizotinib sensitivity of a set of *ALK* mutation-free NSCLC cell lines, especially those with high levels of c-*MET* expression. Notably, chidamide could not increase the crizotinib sensitivity of NSCLC cells cultured in serum-free medium without hepatocyte growth factor (HGF; a c-MET ligand). In contrast, the addition of HGF into the serum-/HGF-free medium could restore the synergistic effect of chidamide. Moreover, the synergistic effect of chidamide could also be abolished either by treatment with c-MET antibody or siRNA-knockdown of c-*MET* expression. While cells with low or no c-*MET* expression were primarily resistant to chidamide-crizotinib cotreatment, enforced c-*MET* overexpression could increase the sensitivity of these cells to chidamide-crizotinib cotreatment. Furthermore, chidamide could decrease c-*MET* expression by inhibiting mRNA N6-methyladenosine (m6A) modification through the downregulation of *METTL3* and *WTAP* expression. Chidamide-crizotinib cotreatment significantly suppressed the activity of c-MET downstream molecules.

**Conclusion:** chidamide downregulated c-*MET* expression by decreasing its mRNA m6A methylation, subsequently increasing the crizotinib sensitivity of NSCLC cells in a c-MET-/HGF-dependent manner.

**GRAPHIC SUMMARY:** 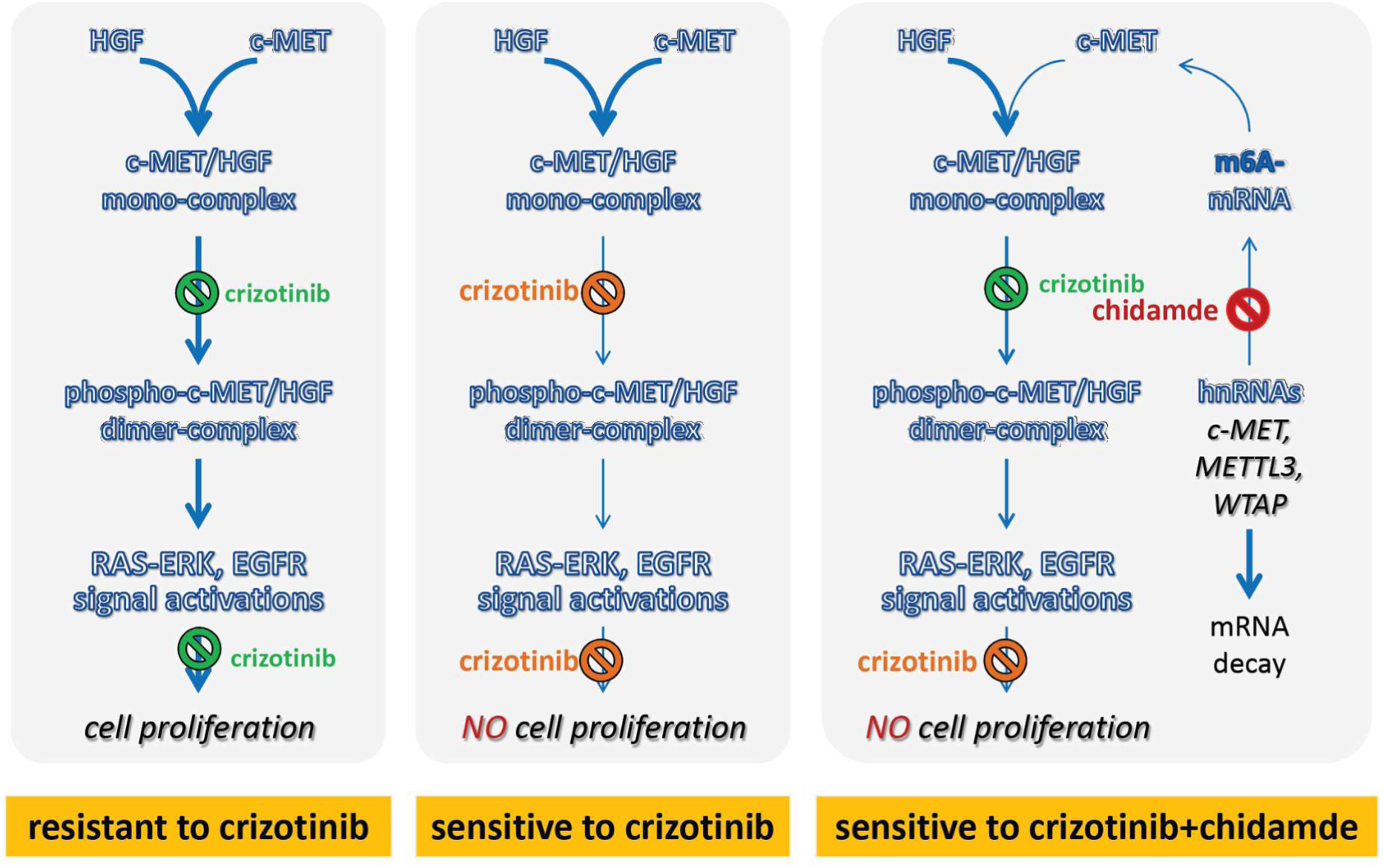

## INTRODUCTION

Non-small cell lung cancer (NSCLC) is the leading cause of cancer death worldwide [1]. Although the recent development of a series of molecular targeted drugs has improved the quality of life and survival of NSCLC patients, a large proportion of NSCLC patients could not benefit from these targeted drugs due to resistance. Therefore, drugs that can reverse primary and secondary drug resistance are urgently expected.

Crizotinib is an ATP-competitive inhibitor targeting ALK/ROS1/c-MET kinases that has been approved by the FDA as a first-line chemical for the treatment of NSCLC and other cancers with *ALK* rearrangement, *ROS1* rearrangement, or aberrant activation of c-*MET* [2-5]. However, the incidence of these alterations in NSCLCs is relatively low (*ALK*, 3-7%; *ROS1*, 1-2%; c-*MET*, 0.8-4%) [4-6], and the majority of NSCLC patients are primarily resistant to treatment with crizotinib [7].

Histone deacetylase inhibitors (HDACIs) can increase the acetylation levels of histone and nonhistone proteins, consequently modulating cancer cell proliferation, apoptosis, differentiation, migration, the host immune response and angiogenesis [8, 9]. Several HDACIs, including vorinostat, panobinostat, and chidamide, have been approved for the treatment of cutaneous T-cell lymphoma, multiple myeloma, and peripheral T-cell lymphoma [10-12]. However, therapeutic effect of HDACIs as single agent is very limited for patients with solid tumors.

Fortunately, increasing evidence suggests that HDACIs combined with other agents show a synergetic increase in antitumor activities [13]. For example, HDACIs could sensitize cancer cells to DNA-damaging drugs by the epigenetic activation of *Schlafen*-11 (*SLFN11*) [14]. The HDACI FA17 could restore the response of breast cancer cells to methotrexate by inhibiting the activity of nucleophosmin and drug efflux pumps through the *PI3K/AKT* pathway [15]. The HDACI LAQ824 could downregulate *BCR-ABL* and sensitize imatinib (an ABL kinase inhibitor) in chronic myelogenous leukemia-blast crisis cells [16]. Several studies have also suggested that HDACIs could enhance the effect of EGFR inhibitors in NSCLC by repressing the expression or phosphorylation of EGFR, HER2, c-MET, AXL, and IGF1R [17-19]. Combinations of HDAC6/8 inhibitors with crizotinib could efficiently inhibit diffuse large B-cell lymphoma and neuroblastoma cells [20][21]. These phenomena suggest that HDACIs could sensitize cancers to different types of drugs and have good application prospects.

Chidamide is a novel HDACI targeting HDAC1/2/3/10 [22]. In this study, we reported for the first time that chidamide could increase the sensitivity of NSCLC cells to crizotinib in a *c-MET* expression-dependent manner *in vitro* and *in vivo*. Our further studies unveiled that chidamide could decrease *c-MET* expression, probably via the downregulation of the RNA methyltransferase *WTAP* and *METTL3* expression and the subsequent loss of *c-MET* m6A mRNA.

## MATERIALS AND METHODS

### Cell lines and culture

In this study, thirteen NSCLC cell lines without *ALK* mutations were used (Table 1). H1299 cells were kindly provided by professor Chengchao Shou, and A549 cells (with KRAS mutations) were kindly provided by professor Zhiqian Zhang. EBC-1 cell line with *c-MET* gene amplification (kindly provided by Dr. Yue Yang) was used as a crizotinib-sensitive control [23]. These two cell lines were tested and authenticated by Beijing JianLian Genes Technology Co., Ltd. before they were used in this study. STR patterns were analyzed using the Goldeneye 20A STR Identifier PCR Amplification Kit. Gene Mapper v3.2 software (ABI) was used to match the STR pattern with those in the online databases of the American Type Culture Collection (ATCC). The other ten cell lines (HCC827, Calu-3, H661, H596, H358, H460, H1650, H1975, H1395, and H292) were purchased from the National Laboratory Cell Resource Sharing Platform (Beijing, China) at the beginning of this study with STR authentications.

**Table 1.** The statuses of related gene mutations^*^ and IC50 values (μM)^**^ of chidamide, crizotinib for 13 NSCLC cell lines with or without chidamide co-treatment.

| Cell line | *ALK*<br>mutation | *ROS1*<br>mutation | *EGFR*<br>mutation | *K-RAS*<br>mutation | *c-MET*<br>amplif. | chidamide<br>IC50 (A) | crizotinib<br>IC50 (B) | crizotinib<br>IC50*** (C) | IC50 Fold<br>change<br>(C to B) | t-test,<br>*p*-value<br>(B vs. C) |
| --- | --- | --- | --- | --- | --- | --- | --- | --- | --- | --- |
| EBC-1 | free | free | free | free | positive | 1.24±0.37 | 0.004±0.00 | 0.003±0.00 | 0.75±0.10 | **0.011** |
| HCC827 | free | free | positive | free | free | 0.90±0.08 | 0.93±0.21 | 0.49±0.09 | 0.54±0.07 | **0.007** |
| H661 | free | free | free | free | free | 5.51±0.77 | 1.10±0.45 | 0.62±0.22 | 0.58±0.06 | **0.006** |
| Calu-3 | free | free | free | free | free | 0.79±0.10 | 4.22±2.22 | 1.98±1.42 | 0.44±0.08 | **0.007** |
| H1299 | free | free | free | free | free | 4.09±0.16 | 1.34±0.13 | 1.10±0.17 | 0.82±0.05 | **0.026** |
| H358 | free | free | free | positive | free | 1.73±0.32 | 1.46±0.61 | 1.13±0.41 | 0.79±0.07 | **0.041** |
| H596 | free | free | free | free | free | 12.6±0.05 | 4.98±1.56 | 4.03±0.92 | 0.81±0.07 | **0.044** |
| H460 | free | free | free | positive | free | 5.99±0.65 | 1.84±0.26 | 1.59±0.23 | 0.86±0.00 | **0.001** |
| A549 | free | free | free | positive | free | 8.25±1.41 | 3.21±1.51 | 3.50±1.37 | 1.11±0.09 | 0.166 |
| H1975 | free | free | positive | free | free | 1.85±0.06 | 1.86±0.66 | 1.94±0.88 | 1.03±0.11 | 0.706 |
| H1650 | free | free | positive | free | free | 92.5±15.7 | 2.17±0.57 | 2.25±0.61 | 1.03±0.03 | 0.166 |
| H292 | free | free | free | free | free | 5.46±0.30 | 1.24±0.11 | 1.66±0.28 | 1.33±0.13 | 0.048 |
| H1395 | free | free | free | free | free | 1.05±0.01 | 4.91±2.29 | 4.85±2.26 | 0.99±0.01 | 0.140 |

\*According to CCLE databases [27]; \*\*Mean ± SD; \*\*\*Cotreated with chidamide at 1/4 IC50

Calu-3 cells were cultured in MEM with 1% nonessential amino acids, EBC-1 cells were cultured in MEM with 1% nonessential amino acids and 1% sodium pyruvate (100mM), H596 cells were cultured in DMEM, and the other ten cell lines were cultured in RPMI 1640 medium. These cell lines, cultured at 37 °C in a humidified incubator with 5% CO_2_, were all supplemented with 10% fetal bovine serum (FBS) and 100 U/mL penicillin/streptomycin (Invitrogen, CA, USA).

### Reagents

Crizotinib (CAS No. 877399-52-5) was purchased from Selleck Chemicals (S1068; Houston, TX, USA). Chidamide (CAS No. 743420-02-2) was kindly provided by Shenzhen Chipscreen Biosciences Ltd. (Shenzhen, China). Human hepatocyte growth factor (HGF; Entrez Gene 3082) was purchased from Sino Biological, Inc. (10463-HNAS, Beijing, China).

### Cell viability assay

Cells were seeded into 96-well plates (3-10×10^3^ cells per well according to different cell growth rates, 5 wells per group) and incubated overnight. Twenty-four hours postseeding, the cells were treated with crizotinib at various concentrations alone or combined with chidamide at an inhibitory concentration of 1/4 IC50. After treatment for 72 hr, the proliferation status of these cells was dynamically determined with a long-term observation platform (IncuCyte, Essen, MI, USA). The cell confluence was analyzed using IncuCyte ZOOM software. Cell viability percent (%) = (cell confluence percent of experimental group/cell confluence percent of control group) × 100%. The half maximal inhibitory concentration (IC50) values were calculated by GraphPad Prism 6 (GraphPad Software, Inc., San Diego, USA). The combination index (CI) values were calculated by CompuSyn version 2.0 software (Biosoft, Cambridge, UK). CI < 1.0 indicates a synergistic effect; CI = 1.0 indicates an additive effect; and CI > 1.0 indicates an antagonistic effect of two drugs [24]. All experiments were conducted three times independently. The IC50 fold change was the ratio of crizotinib-IC50 with and without chidamide and was used to represent the synergistic effect between chidamide and crizotinib.

### Cell apoptosis assay

Approximately 2×10^6^ cells were plated in 10-cm dishes. After the cells attached, they were exposed to DMSO, crizotinib, chidamide or the combination of the two drugs for 48 hr. Then, the cell suspension was collected and incubated with Annexin V-FITC and PI in the dark for 15 min according to the protocol of the Annexin V-Fluorescein Isothiocyanate (FITC) Apoptosis Kit (Dojindo, Kumamoto, Japan). These samples were analyzed by flow cytometry (BD Biosciences, NJ, USA) within 1 hr.

### Cell colony assay

Cells were seeded in 6-well plates at a density of 10^3^ cells per well. After the cells attached, they were treated with DMSO, crizotinib, chidamide or the combination of the two drugs. The medium containing drugs was replaced twice a week for 10-20 days. Finally, the cells were fixed with methyl alcohol and stained with 0.1% crystal violet.

### RNA extraction and quantitative RT-PCR (qRT-PCR) assay

Total RNA was extracted using an Ultrapure RNA kit (Beijing Com Win Biotech Co., China) and reverse-transcribed into cDNA using the First-Strand cDNA Synthesis Kit (Transgen Co., Beijing, China). Then, the cDNA samples were amplified by qRT-PCR using SYBR Green PCR master mix reagents (Roche, Mannheim, Germany) in triplicate. The relative expression levels of the detected genes were normalized to that of the *GAPDH* reference RNA calculated by the classical ΔΔCt method. The sequences (5’-3’) of the primers used are as follows: *c-MET* (Entrez Gene 4233; forward, ccaccctttgttcagtgtgg; and reverse, agtcaaggtgcagctctcat), *ALK* (Entrez Gene 238; forward, gcctgtggctgtcagtatttg; and reverse, tcccatagcagcactccaaag), *ROS1* (Entrez Gene 6098; forward, aggctgccaacatgtctgat; and reverse, cggccagatggtacaggaag), *WTAP* (Entrez Gene 9589; forward, taaagcaacaacagcaggag; and reverse, aatagtccgacgccatca), *METTL3* (Entrez Gene 56339; forward, agtgacagcccagtgcctac; and reverse, acagtccctgctacctccc), *GAPDH* (Entrez Gene 2597; forward, gagatggtgatgggatttc; and reverse, gaaggtgaaggtcggagt), and *18S rRNA* (forward, gagatggtgatgggatttc; and reverse, gaaggtgaaggtcggagt).

### Western blotting

The protein lysates from treated cells were run on an 8%-12% SDS-PAGE gel and transferred onto a PVDF membrane. Then, the membrane was blocked with 5% fat-free milk overnight at 4 °C. The next day, the membrane was incubated with the primary antibodies (MET(D-4)/sc-514148, p-MET(F-5)/sc-377548, STAT3(F-2)/sc-8019, p-STAT3(B-7)/sc-8059, Santa Cruz, USA; AKT(pan) (C67E7)/#4691, p-AKT/#406, ERK(1/2) (137F5)/#4695, p-ERK(Thr202/Tyr204) (D13.14.4E)/#4370, WTAP/#5650, METTL3 (D2I6O)/#96391, METTL14(D8K8W)/#51104, EGFR/#2232, pEGFR(Y1068)/#2234, Cell Signaling Technology, USA; FTO/ab126605, Abcam, UK; and GAPDH/60004-1, Protein Tech, China) at room temperature for at least 1 hr. Then, the membrane was washed with PBST (PBS with 0.1% Tween 20) three times at an interval of 10 min. After washing, the membrane was incubated with the appropriate goat anti-rabbit (SE131, Solarbio, China) or goat anti-mouse (SE131, Solarbio, China) secondary antibodies at room temperature for 1 hr. After washing 6 times, the signals were visualized using the Immobilon Western Chemiluminescent HRP Substrate Kit (WBKLS0500, Millipore, Billerica, USA).

### Plasmid and siRNA transfection

The pLenti-MetGFP vector was kindly provided by David Rimm (Addgene plasmid # 37560; http://n2t.net/addgene: 37560; RRID: Addgene_37560) [25]. The empty vector was constructed by deleting the targeted gene from the pLenti-MetGFP vector. HEK293FT cells were seeded in 6-cm plates before transfection, and when the cell confluence reached 40%, they were transfected with the pLenti-MetGFP vector or the control vector using the lentiviral packaging kit (BG20401S, Beijing Syngentech Co., China) according to the manufacturer’s manual. Then, the cell supernatant, filtered by a 0.45-μm needle filter, was collected and used to infect A549 and H1650 cells 48 hr posttransfection. Afterward, the infected cells were cultured in medium containing 1 μg/mL puromycin (Sigma, St. Louis, USA) for at least two weeks to obtain stably transfected cells.

The siRNA oligo sequences (5’-3’) against c-*MET* mRNAs (#1: sense, gguguugucucaauaucaatt; antisense, uugauauugagacaacacctt; #2: sense, gcaacagcugaaucugcaatt; antisense, uugcagauucagcuguugctt), *WTAP* (#1: sense, cacagaucuuaacucuaautt; antisense, auuagaguuaagaucugugtt; #2: sense, gacccagcgaucaacuugutt; antisense, acaaguugaucgcuggguctt), *METTL3* (#1: sense, gcacauccuacucuuguaatt; antisense, uuacaagaguaggaugugctt; #2: sense, gacugcucuuuccuuaauatt; antisense, uauuaaggaaagagcaguctt) were synthesized by GenePharma Co. (Shanghai, China). HCC827 and H661 cells were transfected with the siRNAs using X-tremeGENETM siRNA Transfection Reagent (Roche, Mannheim, Germany) according to the manufacturer’s manual when the cells reached a confluence of 60-80%. The successful knockdown of c-*MET* expression was confirmed by Western blotting and qRT-PCR. Scramble siRNAs (sense, 5’-uucuccgaacgugucacgutt-3’; antisense, 5’-acgugacacguucggagaatt-3’) were used as the negative control. Cells were further used in other experiments 48 hr posttransfection.

### RNA *N*6-methyladenosine (m6A) dot blotting

Total RNA was extracted from cells in a 6-well microplate (one well per group) using the RNA isolation kit mentioned above. Then, equal amounts of RNA samples were denatured by formaldehyde and spotted on a nylon membrane. After the UV cross-linking of 2500J, the membrane was blocked with a 1% blocking solution (Cat. 11096176001, Roche) for 1 hr. Then, the membrane was incubated with anti-m6A antibody (ab151230, Abcam, UK) at room temperature for at least 2 hr. After washing with 1% blocking solution three times at an interval of 10 min, the membrane was incubated with anti-rabbit secondary antibody (same as above) at room temperature for 1 hr. The input control was dyed by methylthioninium chloride immediately after UV cross-linking.

### m6A methylated RNA immunoprecipitation assay (MeRIP)

Cells in four 10-cm dishes were collected after CHI treatment for 4 hr. The MeRIP assay was carried out using the RNA-Binding Protein Immunoprecipitation Kit (Cat# 17-701, EZ-Magna, Millipore, USA) according to the manufacturer’s instructions. Total m6A-containing RNAs were immunoprecipitated and extracted using m6A antibody (the same as above for m6A dot blotting). cDNA was synthesized from the RIP-RNAs using random primers, and gene-specific quantitative PCR was then performed using the c-*MET* or *18S* RNA control primers in triplicate. The relative c-*MET* mRNA copy number was calculated using that of the *18S rRNA* as an internal reference.

### Animal experiments

HCC827 (3×10^6^) cells suspended in 100 μL of PBS were injected into the left inguen of female Balb/c nude mice (body weight 18-20 g; age 6 weeks; Beijing Huafukang Bioscience Co., Inc.). When the tumor volumes reached 50-100 mm^3^ on the 10th posttransplantation day, the mice were randomized into four groups (10 mice per group) and were intragastrically administered vehicle (normal saline), crizotinib (25 mg/kg body weight), chidamide (5 mg/kg), or the combination of the two drugs daily for 21 days. The tumor volumes and body weights of the mice were measured every 3 days. Tumor volumes were calculated using the following formula: tumor volume (mm^3^) = [(length) × (width)^2^] / 2. These mice were sacrificed by dislocation of cervical vertebra on the 21st treatment day. The CI value in this animal experiment was calculated as follows: CI = [(cA + cB) – cA × cB] / cAB. cA represents the inhibitory rate of chidamide alone, cB represents the inhibitory rate of crizotinib alone, and cAB represents the inhibitory rate of the combination treatment of chidamide and crizotinib. Synergy is defined as a CI value lower than 1.0 [26].

All experiments were performed in accordance with institutional standard guidelines of Peking University Cancer Hospital and adhered to the U.S. Public Health Service Policy on the Humane Care and Use of Laboratory Animals.

### Statistical analysis

SPSS 20.0 software (Armonk, NY, USA) was used to perform the statistical analysis. The Kolmogorov–Smirnov test was used to estimate the normality of distributions. The Mann–Whitney U test was conducted for nonnormally distributed data. Student’s t-test was conducted for normally distributed data. All statistical tests were two-sided. *P* < 0.05 was considered statistically significant.

## RESULTS

### Increasing the sensitivity of NSCLC cells to crizotinib by chidamide

Twelve primary crizotinib-resistant NSCLC cell lines (HCC827, H661, Calu-3, H1299, H460, H596, H358, H292, H1650, H1975, H1395, and A549) with the wild-type *ALK* gene were selected according to Pharmacogenomics datasets of the Cancer Cell Line Encyclopedia (CCLE) project [27] and used in this study. First, we examined the cytotoxicity and IC50 value of chidamide as a single agent for these cell lines and found that the effect of chidamide was time- and dose-dependent (range: 0.79 - 92.5 µM, Table 1; Figure S1A-L). Next, we investigated the antiproliferative activities of crizotinib at various concentrations with and without chidamide at ≤1/4 IC50 (final concentration, ≤ 1.0 µM) for each cell line. Compared to crizotinib treatment alone, we found that chidamide-crizotinib cotreatment decreased viability in seven cell lines (HCC827, H661, Calu-3, H1299, H460, H596, and H358; Figure 1A-G, top charts). In these cells, the chidamide-crizotinib CI values were <1.0, and the crizotinib IC50 values for the chidamide (at 1/4 IC50)-treated cell lines were significantly lower than those treated with crizotinib alone (HCC827, *p* = 0.007; H661, *p* = 0.006; Calu-3, *p* = 0.007; H1299, *p* = 0.026; H358, *p* = 0.041; H596, *p* = 0.044; and H460, *p* < 0.001) (Figure 1A-G, bottom chart). Especially for the chidamide-cotreated HCC827, H661, and Calu-3 cell lines, the CI values were < 0.7, and the crizotinib IC50 values decreased > 50% (Table 1). For the H292, H1650, H1975, H1395, and A549 cell lines, the synergistic effect between crizotinib and chidamide was not seen, and the CI values were ≥ 1 (Figure 1H-L). An antagonistic effect was detected in H292 cells (*p* = 0.048). These data revealed that chidamide could synergistically enhance the cytotoxic effects of crizotinib in 7 of the 12 NSCLC cell lines that are not crizotinib sensitive.

**Figure 1.**
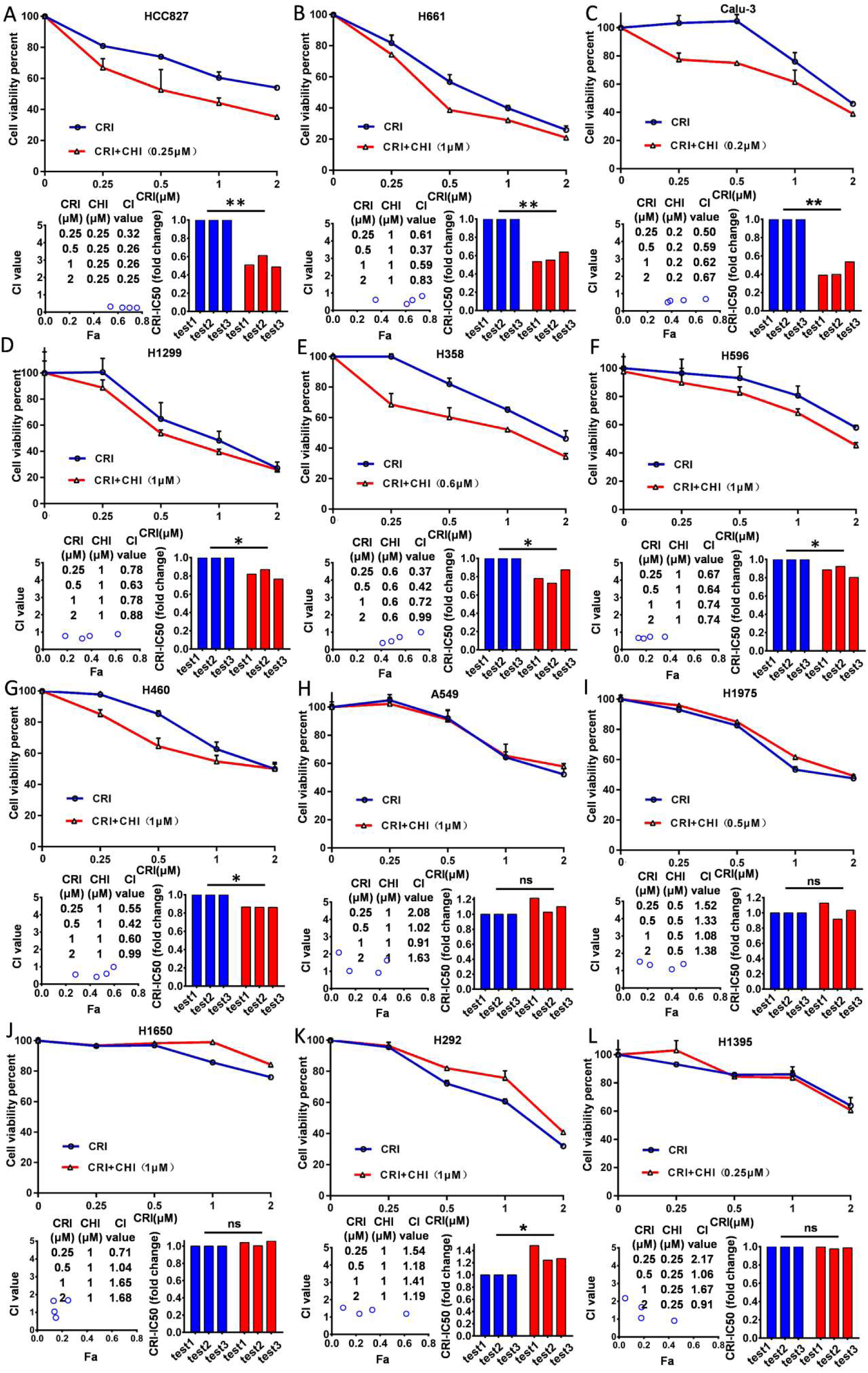
Effect of chidamide on the sensitivity of ALK mutation-free NSCLC cells to crizotinib. NSCLC cells were treated with various concentrations of crizotinib (CRI) alone or combined with chidamide (CHI) at ≤1/4 IC50 for 72 hr. Cell viability was measured by using the IncuCyte platform (top charts). The crizotinib-IC50 values for 12 NSCLC cell lines were calculated in the absence or presence of chidamide, and the fold change of crizotinib-IC50 in three independent experiments is presented as a histogram (bottom right charts). The synergetic effect of chidamide-crizotinib cotreatment on cell proliferation inhibition was calculated using the CI equation and presented as Fa (fraction affected by the dose) in the Fa–CI plots (bottom left charts). */**/***, *p*<0.05/0.0/0.001; ns, not significant; (**A**) HCC827; (**B**) H661; (**C**) Calu-3; (**D**) H1299; (**E**) H358; (**F**) H596; (**G**) H460; (**H**) A549; (**I**) H1975; (**J**) H1650; (**K**) H292; (**L**) H1395.

To further validate the chidamide-crizotinib synergistic effect, cell apoptosis and colony formation assays were performed using the HCC827, Calu-3, and H661 cell lines. Compared with single drug treatment, chidamide-crizotinib cotreatment significantly increased the percentage of apoptotic cells (67.7% for HCC827; 58.7% for H661; and 34.6% for Calu-3; *p* < 0.050; Figure 2A). Chidamide-crizotinib cotreatment also significantly inhibited colony formation in all of these cell lines (6.7% for HCC827; 3.9% for H661; and 8.6% for Calu-3 cells; *p* < 0.050; Figure 2B).

**Figure 2.**
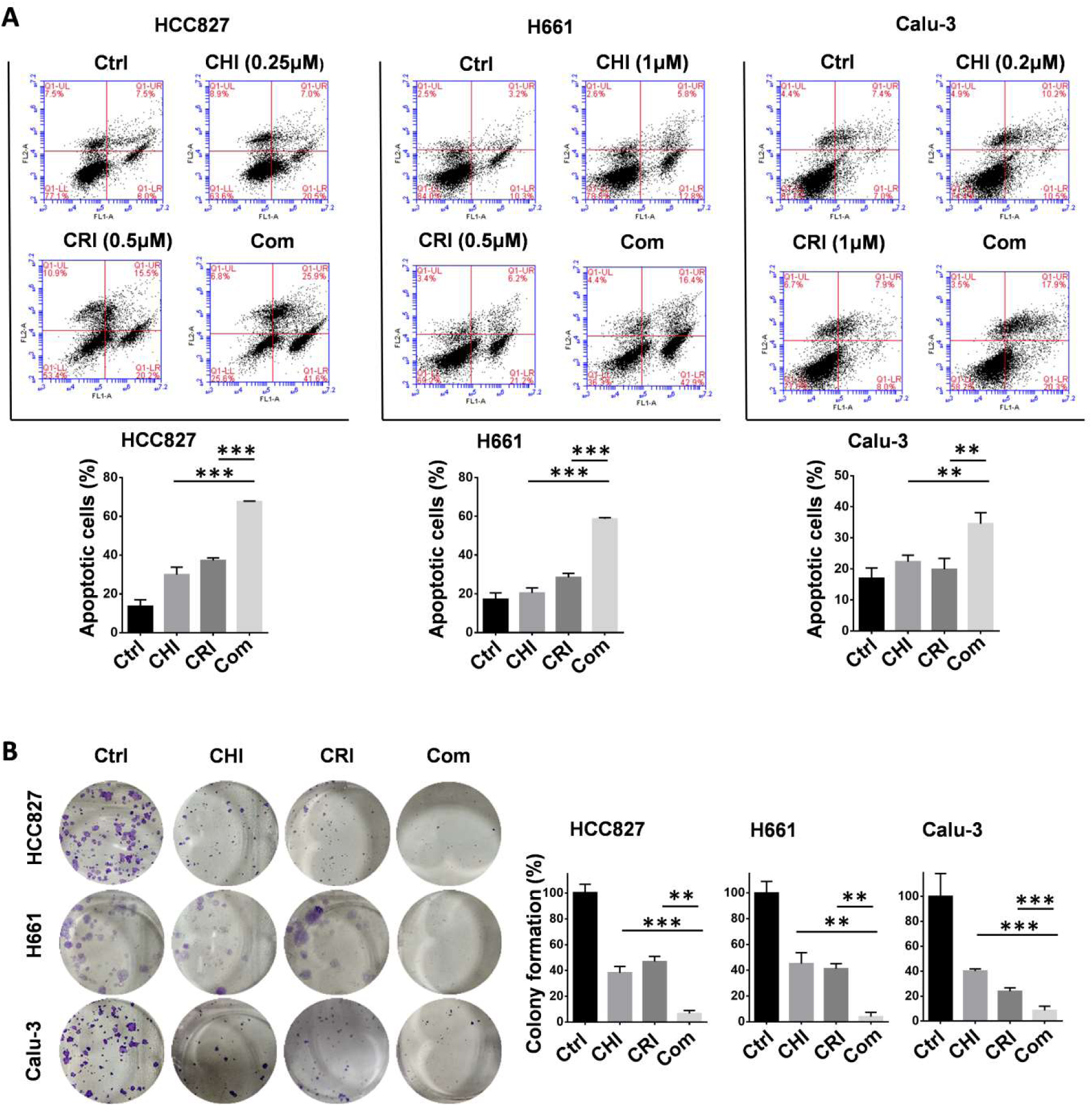
Effect of chidamide on the crizotinib-induced apoptosis and growth arrest of NSCLC cells. (**A**) Effect of chidamide (CHI), crizotinib (CRI), and chidamide-crizotinib cotreatment on the apoptosis of HCC827, H661, and Calu-3 cells. Cells were treated with chidamide (at 1/4 IC50) and crizotinib (0.5 or 1 μM) alone or in combination for 48 hr. DMSO was used as an agent control (Ctrl). Apoptotic cells were labeled with propidium iodide/Annexin V-FITC and analyzed by flow cytometry. (**B**) Effect of chidamide, crizotinib, and cotreatment on the colony formation of HCC827, H661, and Calu-3 cells. These cells were treated with the indicated concentrations of chidamide or crizotinib for 10–20 days. Colonies were stained with 0.1% crystal violet and photographed (left images). The results represent the mean of three independent experiments (right charts). **/***, *p*<0.01/0.001.

Overall, these data suggested that chidamide-crizotinib cotreatment apparently promoted the cell apoptosis and inhibited the cell proliferation of NSCLC cells compared with single drug treatment *in vitro*.

### Enhancement of crizotinib sensitivity by chidamide in a c-MET expression-dependent manner

To explore the underlying mechanism of enhancing the sensitivity of NSCLC cells to crizotinib by chidamide, the baseline mRNA levels of the crizotinib target genes *ALK, ROS1* and c-*MET* were determined. We found that in cell lines in which chidamide could enhance the sensitivity of crizotinib, the level of c-*MET* expression was much higher than that in cell lines in which chidamide could not increase the sensitivity of crizotinib (Figure 3A). The level of c-*MET* mRNA was inversely correlated with the chidamide-based crizotinib-IC50 fold change (the smaller the change value was, the stronger the sensitization effect) (*n* = 12, *r* = −0.706, *p* = 0.010; Figure 3B; Table 1). No significant association was observed between the chidamide-based crizotinib-IC50 fold change and the level of *ALK* or *ROS1* mRNA. The Western blotting results confirmed the qRT-PCR results (Figure 3C). Amounts of the c-MET protein were also inversely correlated with the chidamide-based crizotinib-IC50 fold change (*n* = 12, *r* = −0.694, *p* = 0.012). No significant relationship between the level of the c-MET protein and crizotinib-IC50 or chidamide-IC50 was found (Figure 3D). These phenomena imply the possibility that the synergistic effect of chidamide on crizotinib sensitivity in NSCLC cells may depend on the level of c-*MET* expression.

**Figure 3.**
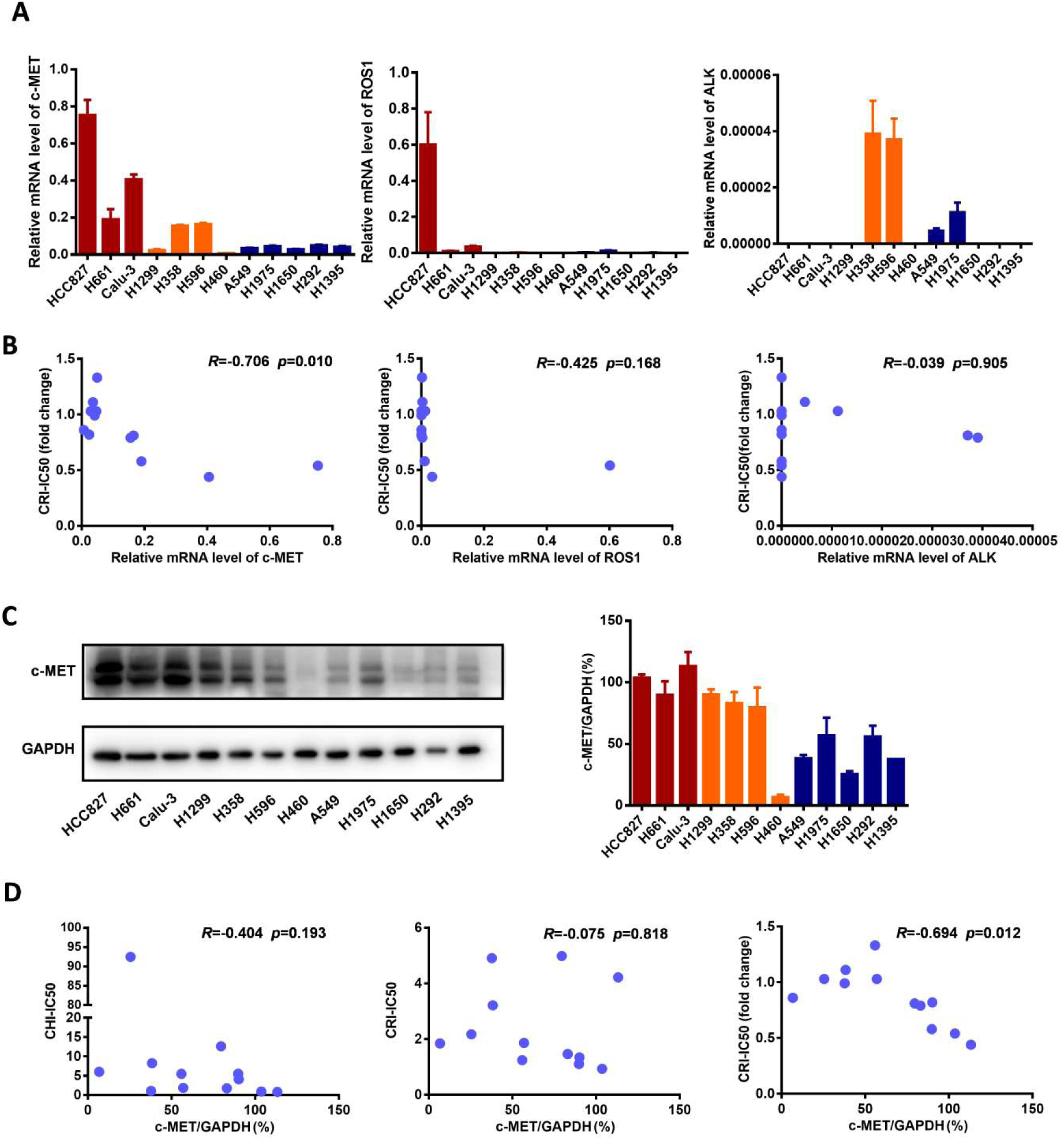
Comparisons of c-*MET, ROS1, ALK* expression levels in NSCLC cells with different crizotinib sensitivities. (**A**) The levels of c-*MET, ROS1, ALK* mRNAs in 12 NSCLC cell lines by qRT-PCR. (**B**) Relationships between the crizotinib IC50 fold change and baseline mRNA expression levels of c-*MET, ROS1, ALK* genes in 12 NSCLC cell lines. **(C)** The baseline levels of c-MET protein in 12 NSCLC cell lines by Western blotting. The density of each band was calculated by ImageJ software, and the ratio of c-MET/GAPDH is presented by means with bars representing the SEM. **(D)** Relationships between the levels of c-MET protein and chidamide IC50, crizotinib IC50 (without chidamide), and crizotinib IC50 fold change (with chidamide) in 12 NSCLC cell lines. Pearson correlation analyses were performed to calculate the correlation coefficient (*R*) and *p*-value.

Therefore, we further studied whether c-MET could play a driver role in the synergetic effect of chidamide-crizotinib. In the serum-free medium not containing HGF (a c-MET ligand), no synergetic effect was found in the above chidamide-crizotinib-sensitive cell lines HCC827 and H661 with high baseline *c-MET* expression. However, the synergetic effect was restored with the addition of HGF (final concentration, 100 ng/mL) into the serum-/HGF-free medium (Figure 4A). In addition, the synergetic effect could be reversed in cells either by treatment with c-MET antibody (Figure 4B) or by c-*MET* knockdown (Figure 4C). Loss of function of c-MET by antibody or HGF-deprivation could also inhibit the proliferation of HCC827 and H661 cells with chidamide treatment (Figure S2A). In addition, the HGF supplement itself significantly promoted the proliferation of these cells. In contrast, the c-MET antibody blocking itself significantly inhibited these cell proliferation (Figure S2B). These results suggest that the acitivity of c-MET signal pathway may affect these cell proliferation.

**Figure 4.**
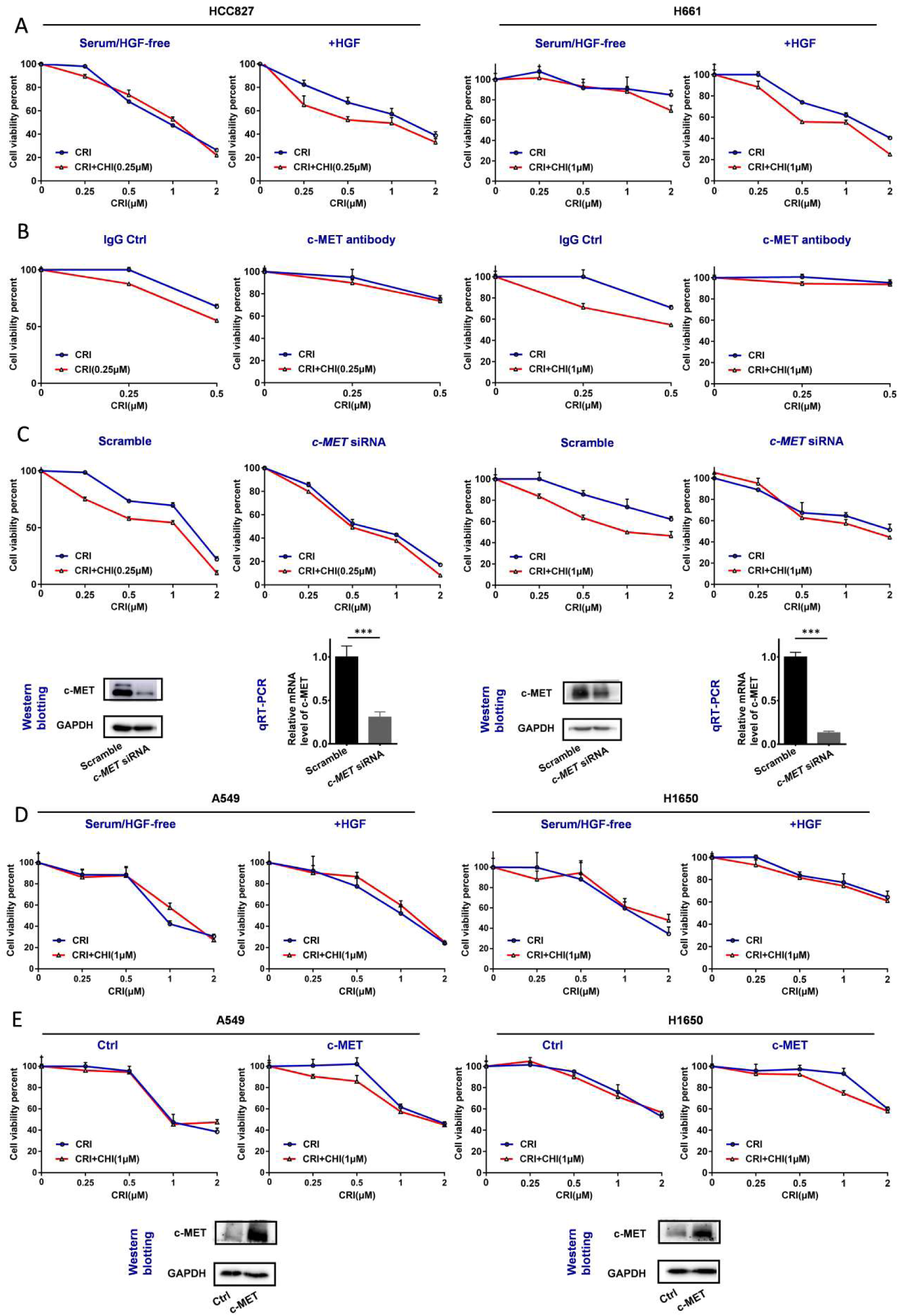
Effect of the loss and gain of c-MET function on increasing the crizotinib sensitivity of NSCLC cells by chidamide. (**A**) The increasing of crizotinib sensitivity by chidamide in two chidamide-crizotinib-sensitive cell lines, HCC827 and H661, cultured in serum-free medium with no c-MET ligand HGF or with 100 ng/mL HGF supplement. (**B**) Effect of c-MET antibody treatment (2.5 μg/mL) on the increasing of crizotinib sensitivity by chidamide in HCC827 and H661 cells. IgG (2.5 μg/mL) was used as a negative antibody control. (**C**) Effect of siRNA knockdown of c-*MET* mRNA on the increasing of crizotinib sensitivity by chidamide in HCC827 and H661 cells. Cells transfected with the scramble siRNA were used as the transfection agent control. (**D**) Comparison of the increasing of crizotinib sensitivity by chidamide in two chidamide-crizotinib non-sensitive control cell lines (A549 and H1650) cultured in serum-free medium with no HGF or with 100 ng/mL HGF supplement. (**E**) Effect of enforced c-*MET* overexpression on the increasing of crizotinib sensitivity of the A549 and H1650 cell lines by chidamide.

For the two chidamide-crizotinib-resistant control cell lines A549 and H1650, the synergetic effect of chidamide-crizotinib was not different between the serum-free medium with and without the addition of HGF (Figure 4D). However, enforced c-*MET* overexpression could induce the synergetic effect of chidamide-crizotinib (Figure 4E). In EBC-1 cells with *c-MET* gene amplification, a significant increase of the level of *c-MET* mRNA, but not the amount of c-MET protein, was detected relative to those of two control cell lines HCC827 and Calu-3 (Figure S3A-B). Interestingly, the synergetic effect of chidamide-crizotinib was also observed in the representative crizotinib-sensitive EBC-1 cells and chidamide could downregulate the level of *c-MET* expression (Figure S3C-D). Overall, the above data indicate that the synergetic effect of chidamide-crizotinib is dependent on c-*MET* expression. Low or no c-*MET* expression leads to the absence of a synergetic effect.

### Downregulation of *c-MET* expression by chidamide via RNA m6A modification

We further studied the mechanisms of chidamide in increasing crizotinib sensitivity. After treatment with chidamide for 4 hr, the amounts of c-MET protein by Western blotting in three chidamide-crizotinib-sensitive cell lines (HCC827, H661 and Calu-3) were significantly decreased while the levels of c-*MET* mRNA by qRT-PCR were not significantly changed (Figure 5A), suggesting a posttranscriptional mechanism.

**Figure 5.**
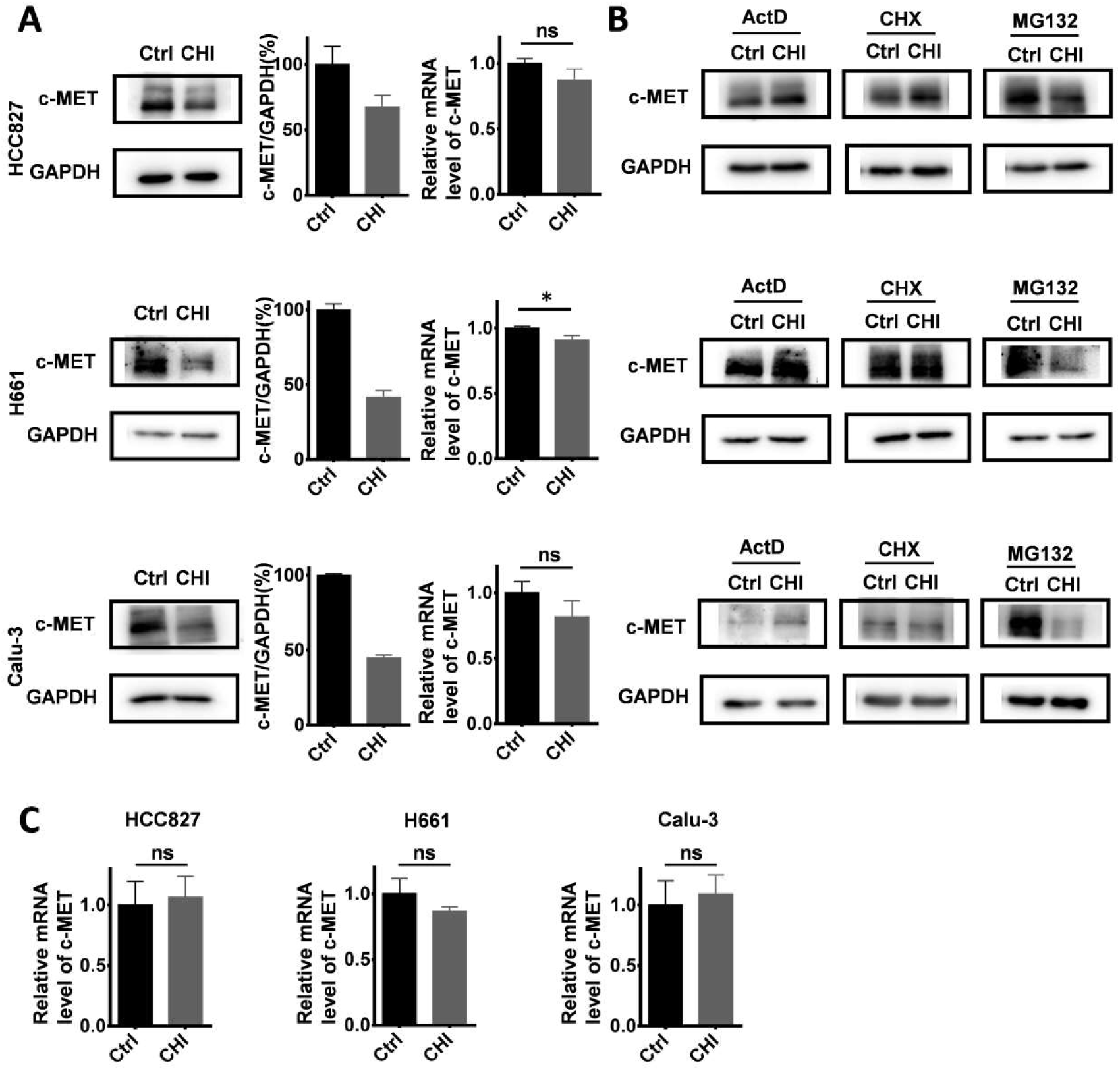
Effect of chidamide on the levels of c-*MET* expression in NSCLC cells. (**A**) Effects of chidamide treatment (0.25 μM for HCC827, 1 μM for H661, 0.2 μM for Calu-3; 4 hr) on the levels of c-*MET* expression in HCC827, H661, and Calu-3 cells in Western blotting (left images and middle charts) and qRT-PCR (right charts) analyses. (**B**) Effects of treatment with transcription, translation, and proteasome inhibitors (ActD, CHX, and MG132) on the chidamide-induced downregulation of c-*MET* expression in HCC827, H661, and Calu-3 cells by Western blotting. These cells were exposed to chidamide, ActD (5 μg/mL), CHX (50 μg/mL), or MG132 (10 μM) alone or cotreated with chidamide and ActD, CHX, or MG132 for 4 hr. (**C**) The level of c-*MET* mRNA in HCC827, H661, and Calu-3 cells cotreated with ActD. *, *p*<0.05; ns, not statistically significant.

To dissect which of the posttranscriptional processes chidamide may affect to influence the amount of c-MET protein, these cell lines were cotreated with several known inhibitors, including actinomycin D (ActD; 5 μg/mL), cycloheximide (CHX; 50 μg/mL), and proteasome inhibitor MG132 (10 μM), for 4 hr to block RNA transcription, protein synthesis, and degradation, respectively (Figure 5B). Western blotting analyses showed that the downregulation of *c-MET* expression by chidamide remained in the cells cotreated with MG132. However, the amount of c-MET protein was not decreased in the cells cotreated with CHX or ActD, and the level of c-*MET* mRNA was unchanged by ActD (Figure 5C). These results suggest that both ubiquitin-related protein degradation and RNA transcription may not be involved in downregulation of c-MET expression by chidamide, and chidamide may decrease the amount of c-MET protein from the posttranscription to translation steps.

RNA modifications regulate RNA splicing, translation, and degradation, and m6A is the main RNA modification [28]. To study whether chidamide affects c-*MET* m6A mRNA modification, we compared the levels of global and c-*MET* m6A mRNA in these cells with and without chidamide treatment. Dot blotting showed that chidamide could markedly decrease the total m6A RNA level of HCC827 and H661 cells (Figure 6A). The results of the MeRIP-based qRT-PCR assay further demonstrated that a significant decrease in the relative c-*MET* m6A copy number was induced in the chidamide-crizotinib-sensitive cell lines HCC827 and H661 by chidamide (−1.83 for HCC827; −1.91 for H661; Figure 6B). Although the total levels of total m6A RNA were also decreased in the two chidamide-crizotinib-resistant control cell lines A549 and H1650 by chidamide (Figure 6A), a very limited decrease in the relative c-*MET* m6A copy number could be induced due to the low baseline level of *c-MET* mRNA in these cells (−0.11 for A549; −0.24 for H1650; Figure 6B). This finding demonstrates that chidamide (at 1/4 IC50) treatment may increase crizotinib sensitibity by inhibiting the m6A modification of c-*MET* mRNA.

**Figure 6.**
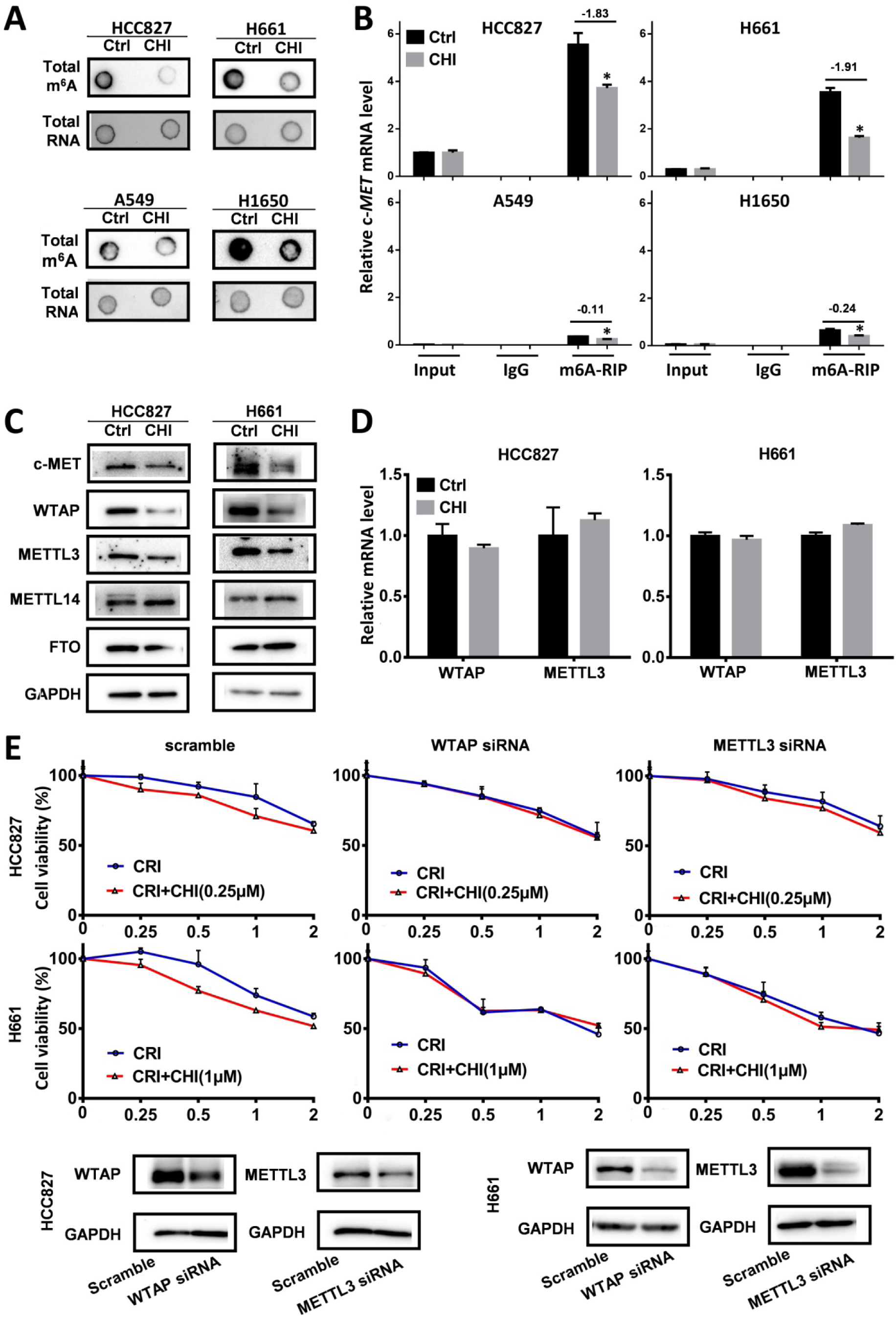
Effect of RNA m6A methylation changes on increasing the crizotinib sensitivity of NSCLC cells by chidamide. (**A**) The amounts of total m6A RNA of two chidamide-crizotinib-sensitive cell lines (HCC827 and H661) and two chidamide-crizotinib-resistant cell lines (A549 and H1650) treated with chidamide (0.25 μM for HCC827; 1 μM for H661, A549, and H1650; 4 hr) in dot blotting analyses. (**B**) Relative copy number (RCN) of c-*MET* m6A mRNA in chidamide-crizotinib-sensitive and -resistant cell lines with and without the treatment of chidamide in MeRIP-based qRT-PCR analyses. The average input RCN value of the c-*MET* mRNA of HCC827 control cells was used to normalize all RCN values of the c-*MET* mRNA of the 4 cell lines in different groups. The RCN difference of c-*MET* m6A mRNA between cells with and without chidamide treatment is labeled. (**C**) Amounts of m6A methylation writer complex components WTAP and METTL3 proteins in HCC827 and H661 cells with and without chidamide treatment in Western blotting. (**D**) The levels of *WTAP* and *METTL3* mRNAs in chidamide-treated HCC827 and H661 cells. (**E**) Effect of siRNA knockdown of *METTL3* or *WTAP* expression on the synergetic effect of chidamide-crizotinib in HCC827 and H661 cells. **/***, *p*<0.01/0.001

To study how chidamide affects RNA m6A modification, the expression levels of several known m6A writer complex components (METTL3, METTL14, and WTAP) and eraser (FTO) were determined. The Western blotting results revealed that the amounts of METTL3 and WTAP proteins were consistently decreased (Figure 6C), and the levels of *METTL3* and *WTAP* mRNAs were not significantly changed in chidamide-treated HCC827 and H661 cells (Figure 6D). In addition, the synergetic effect of chidamide-crizotinib disappeared in both HCC827 and H661 cells with siRNA knockdown of *METTL3* or *WTAP* expression (Figure 6E). These data suggest that the downregulation of *METTL3* and *WTAP* expression may play a crucial role in the increasing of crizotinib sensitivity by chidamide.

### Synergistic inhibition of the growth of NSCLC by chidamide-crizotinib cotreatment in mice

To further evaluate the effect of chidamide-crizotinib cotreatment *in vivo*, we subcutaneously injected HCC827 cells into the left inguen of female Balb/c mice (10 mice per group). When the tumor volumes reached 50-100 mm^3^, these mice with established HCC827-derived tumors were treated (i.g.) with vehicle, chidamide and crizotinib alone or their combination for 21 days. We found that while chidamide or crizotinib treatment alone only weakly decreased the tumor weight compared to the vehicle control, chidamide and crizotinib cotreatment led to a markedly decreased growth of tumors on the 21st treatment day (Figure 7A-B). Chidamide-crizotinib cotreatment exhibited ideal synergistic effects (CI = 0.880). Moreover, chidamide and crizotinib treatment alone or in combination did not lead to a loss of body weight (Figure 7C) or to other apparent illnesses, suggesting high safety. These findings demonstrate that chidamide could increase the sensitivity of NSCLC cells to crizotinib *in vivo*.

**Figure 7.**
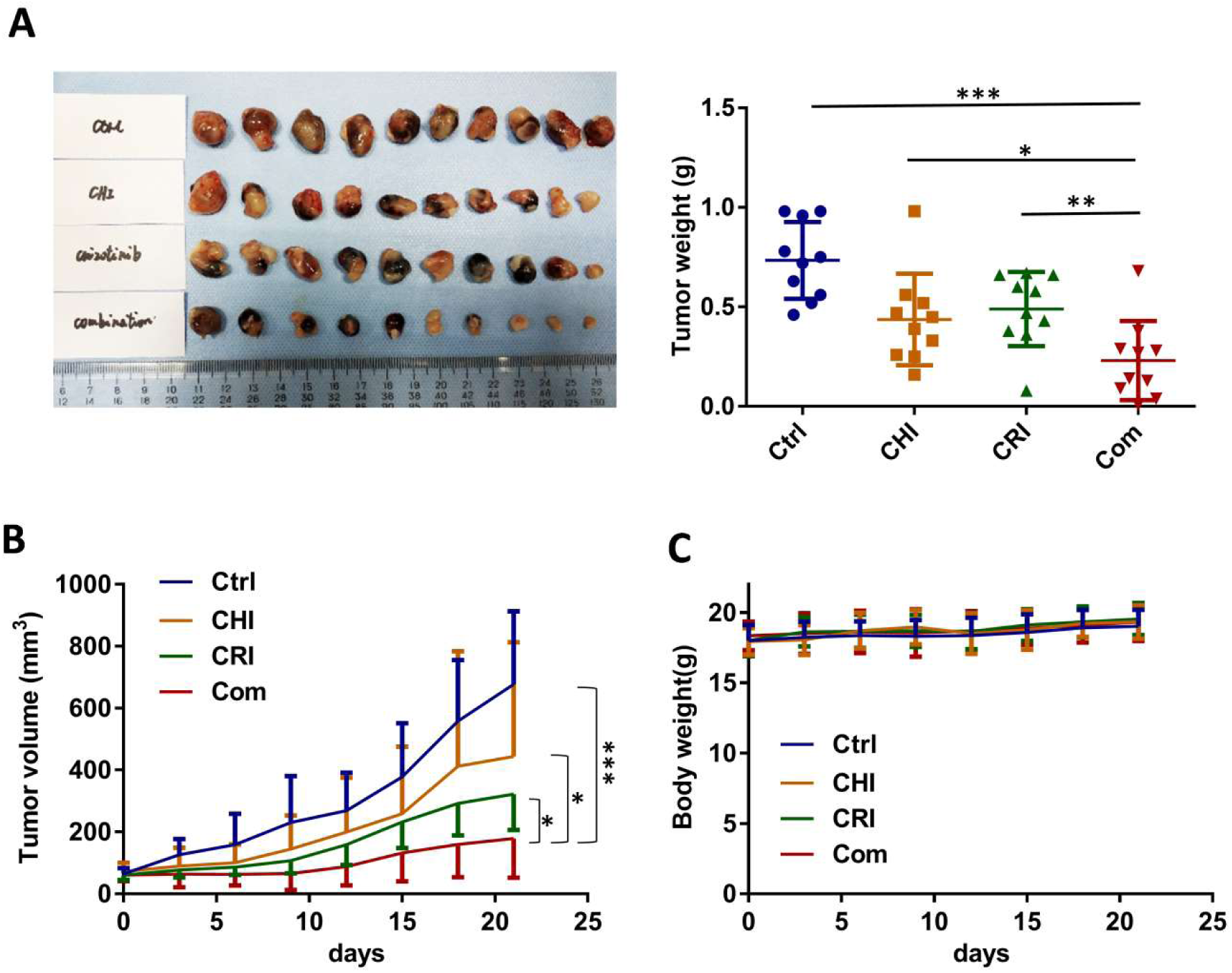
Effect of the combined treatment of chidamide and crizotinib on the growth of HCC827-derived tumor xenografts in Balb/c nude mice. (**A**) The growth status of HCC827 xenografts in mice treated with chidamide, crizotinib, or their combination (n=10) on the 21st day of treatment. Balb/c nude mice bearing established HCC827-derived tumors were treated with vehicle, chidamide (5 mg/kg/d), crizotinib (25 mg/kg/d) or drug combinations by intragastric administration daily for 21 days. (**B**) Tumor volumes in different groups. (**C**) Body weights of mice in different groups. */**/***, P <0.05/0.01/0.001

It was reported that crizotinib inhibited tumor growth by decreasing the phosphorylation levels of c-MET, ERK, AKT, and STAT3 [29, 30]. In HCC827 and H661 cell lines that are not crizotinib-sensitive, crizotinib treatment alone could not affect the phosphorylation levels of ERK and STAT3 proteins in HCC827 cells or the phosphorylation levels of c-MET and ERK proteins in H661 cells. However, the phosphorylation levels of all four of these proteins were consistently decreased in both cell lines with chidamide-crizotinib cotreatment (Figure S4A). The effects of chidamide-crizotinib cotreatment on the phosphorylation of these proteins were also observed in representative mouse HCC827-derived tumors (Figure S4B).

## Discussion

Chidamide is a benzamide HDAC inhibitor approved for the treatment of cutaneous T-cell lymphoma. Studies suggest that chidamide may sensitize some DNA damage or target-specific drugs [10, 31, 32]. Crizotinib is a first-line chemical approved for the treatment of NSCLC and other cancers with *ALK* and *ROS1* mutations or c-*MET* amplification [2-5]. In the present study, we found, for the first time, that chidamide can increase the crizotinib sensitivity of *ALK* mutation-free NSCLC cells in a c-MET/HGF-dependent manner. Furthermore, the downregulation of RNA m6A writer *METTL3* and *WTAP* expression and consequent loss of c-*MET* m6A mRNA is the possible mechanism for chidamide to increase the crizotinib sensitivity of NSCLC cells actively expressing c-*MET*.

The membrane HGF receptor c-MET is a tyrosine kinase that can activate the PI3K-AKT, Ras-MAPK, and STAT3 signaling pathways and promote cancer cell proliferation, migration, and invasion [29, 30, 33]. Recent studies have found that the c-MET-HGF axis also plays an important role in the occurrence and development of NSCLC [34-37]. There are many mechanisms leading to the upregulation of c-*MET* overexpression, including gene copy number amplification and posttranslational modifications [38].

Although c-MET is one of the targets of crizotinib, only patients with c-*MET* amplification or exon-14 skipping mutation are currently approved to receive crizotinib treatment. Relative to the low incidence of c-*MET* amplification, c-*MET* overexpression occurs in 35–72% of NSCLCs [38-42]. In our study, we found that the amount of c-MET protein was not associated with the crizotinib IC50 value in 12 NSCLC cell lines. The mining result of the Cancer Cell Line Encyclopedia (CCLE) dataset [27] confirmed our results that no correlation between the level of c-*MET* mRNA and the crizotinib IC50 value of pan-cancer cell lines (Figure S5A) or lung cancer cell lines could be observed (Figure S5B). While NSCLCs with c-*MET* amplification or exon14 splicing alterations were responsive to crizotinib treatment, some NSCLCs with high c-*MET* expression were primarily resistant to crizotinib, which may account for why c-*MET* overexpression is not a biomarker for treatment with crizotinib [5,43,44]. Interestingly, we found that NSCLC cells overexpressing c-*MET* were sensitive to chidamide-crizotinib cotreatment *in vitro* and *in vivo*. It is useful to study whether chidamide could increase the sensitivity of other kinds of cancer cells to crizotinib treatment.

Previous studies have demonstrated that the inhibition of HDAC could reduce the expression of c*-MET* and the phosphorylation of extracellular signal-regulated kinases 1 and 2 by decreasing the levels of *hsa-miR-449a* in hepatocellular carcinoma cells [45]. The inhibition of HDAC6 could also downregulate c-*MET* expression while promoting c-MET ubiquitination-dependent degradation in human diffuse large B-cell lymphoma [20]. HDACIs, including chidamide and YF454A, could downregulate the expression of c-MET and sensitize the effect of EGFR TKIs on the proliferation of NSCLC cells [17, 32]. HCC827 cells contain mutant EGFR gene, which may lead to them not to be sensitive to crizotinib due to remained activity of mutant EGFR in these cells. It was reported that erotinib, an EGFR inhibitor, could increase the sensitivity of cell to crizotinib [46]. To study whether chidamide might increase the crizotinib sensitivity through downregulation of EGFR phosphorylation, we downregulated EGFR expression in HCC827 cells by siRNA. We found that the synergetic effect of chidamide-crizotinib could not be abolished by the downregulation of EGFR expression (Figure S6A and S6B), although chidamide, crizotinib, and their combination indeed partially decrease the phosphorylation level of EGFR protein in HCC827 cells (Figure S6C) and its derived tumors (Figure S6D). These results suggest that the synergetic effect may not depend on the decrease of EGFR phosphorylation. Instead, the synergetic effect of chidamide-crizotinib should depend on the downregulation of *c-MET* expression.

In the present study, we found that chidamide treatment could decrease the amount of c-MET protein, but not c-*MET* mRNA, in several NSCLC cell lines. Treatment with the proteasome inhibitor MG132 did not block the decrease in c-MET protein in these cells. However, the ribosome inhibitor CHX treatment completely blocked the decrease in c-MET protein by chidamide. These phenomena suggest that chidamide may posttranscriptionally downregulate c-*MET* expression.

RNA m6A modification discovered in the 1970s is a main modification of eukaryotic messenger RNAs (mRNAs), microRNAs (miRNAs), and long noncoding RNAs (lncRNAs) that regulates gene expression via changes in RNA splicing, translation efficiency, and stability [27, 47-49]. Our results showed that chidamide could downregulate the m6A level of total RNA. The results of the MeRIP-based qRT-PCR analyses further demonstrated that the c-*MET* m6A copy number was also significantly decreased by chidamide in chidamide-crizotinib-sensitive cell lines but not in chidamide-crizotinib-resistant cell lines. Further study showed that chidamide could downregulate the levels of m6A methyltransferase METTL3 and WTAP proteins. siRNA knockdown of *METTL3* and *WTAP* expression completely abolished the synergistic effect between chidamide and crizotinib. These results indicate that treatment with chidamide can repress methyltransferase *METTL3* and *WTAP* expression and decrease the c-MET protein level through the loss of c-*MET* mRNA m6A methylation. The underlying mechanism of the downregulation of *METTL3* and *WTAP* expression by chidamide is worth further studying.

In conclusion, the present study demonstrates that chidamide can decrease the level of c-*MET* expression by inhibiting mRNA m6A methylation and can subsequently increase the sensitivity of c-*MET*-expressing NSCLC cells to crizotinib. The combination of chidamide and crizotinib may be a promising novel strategy for the treatment of NSCLC with high levels of c-MET expression or c-*MET* gene amplification. Clinical trials should be conducted to prove our findings.

## Abbreviations

NSCLC: non-small cell lung cancer;
HGF: the hepatocyte growth factor;
HDACI: histone deacetylase inhibitor;
MeRIP: m6A methylated RNA immunoprecipitation assay;
IC50: the half maximal inhibitory concentration;
CI: the combination index;
CCLE: the Cancer Cell Line Encyclopedia;
CHX: cycloheximide;
ActD: actinomycin D.

## Acknowledgements

This work was financially supported by a grant from the Beijing Municipal Commission of Health and Family Planning (PXM2018_026279_000005) to DD. We thanks Dr. Xiangping Lu Shenzhen Chipscreen Biosciences Ltd. (Shenzhen, China) for kindly providing us chidamide.

