## Supplementary Figures 1-6 for "Chidamide increases the sensitivity of non-small cell lung cancer to crizotinib by decreasing c-*MET* mRNA methylation"

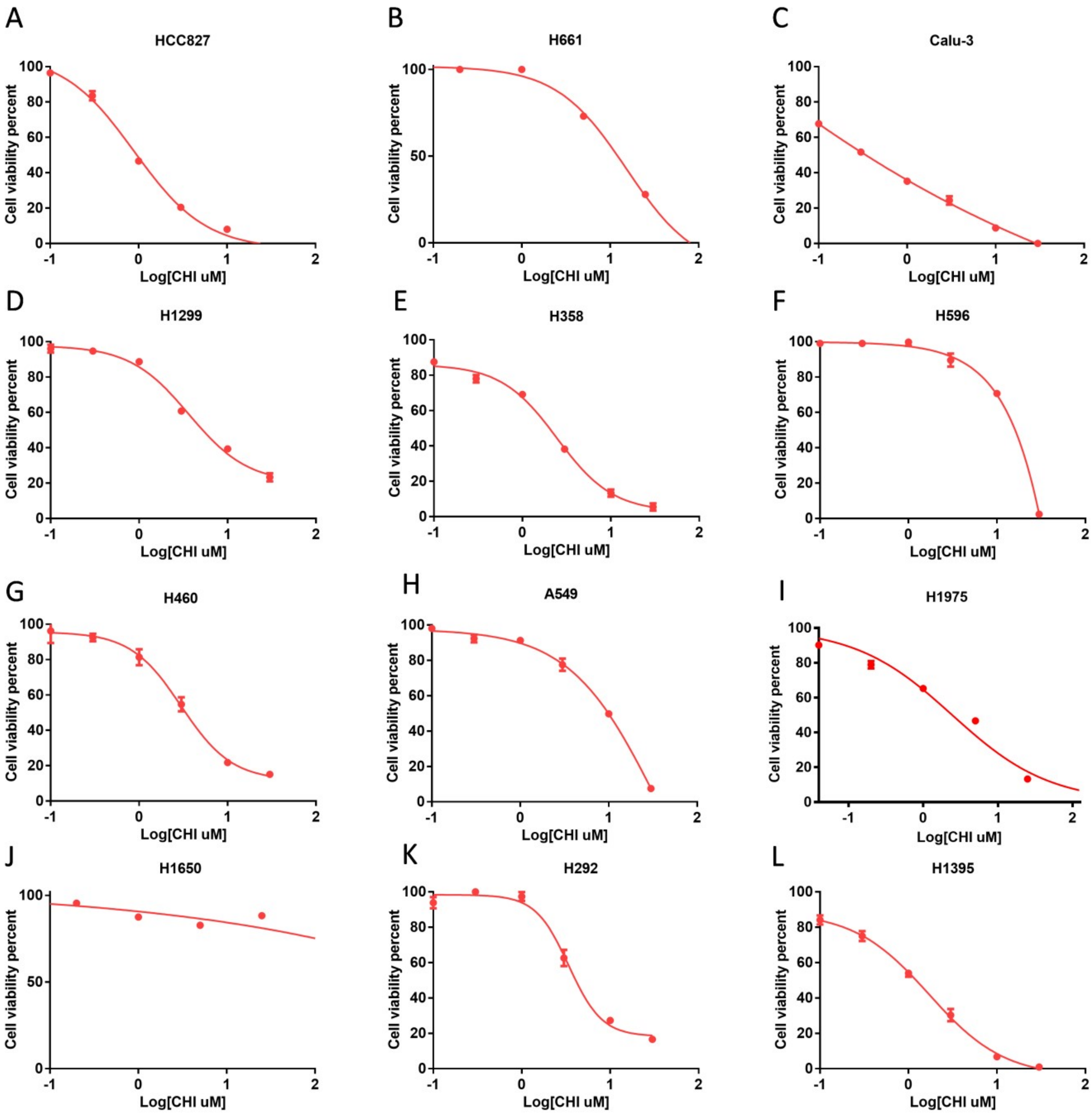


**Figure S1.** Cell viability curves of 12 NSCLC cell lines treated with chidamide (CHI) at various concentrations for 72 hr. (**A**) HCC827; (**B**) H661; (**C**) Calu-3; (**D**) H1299; (**E**) H358; (**F**) H596; (**G**) H460; (**H**) A549; (**I**) H1975; (**J**) H1650; (**K**) H292; (**L**) H1395.

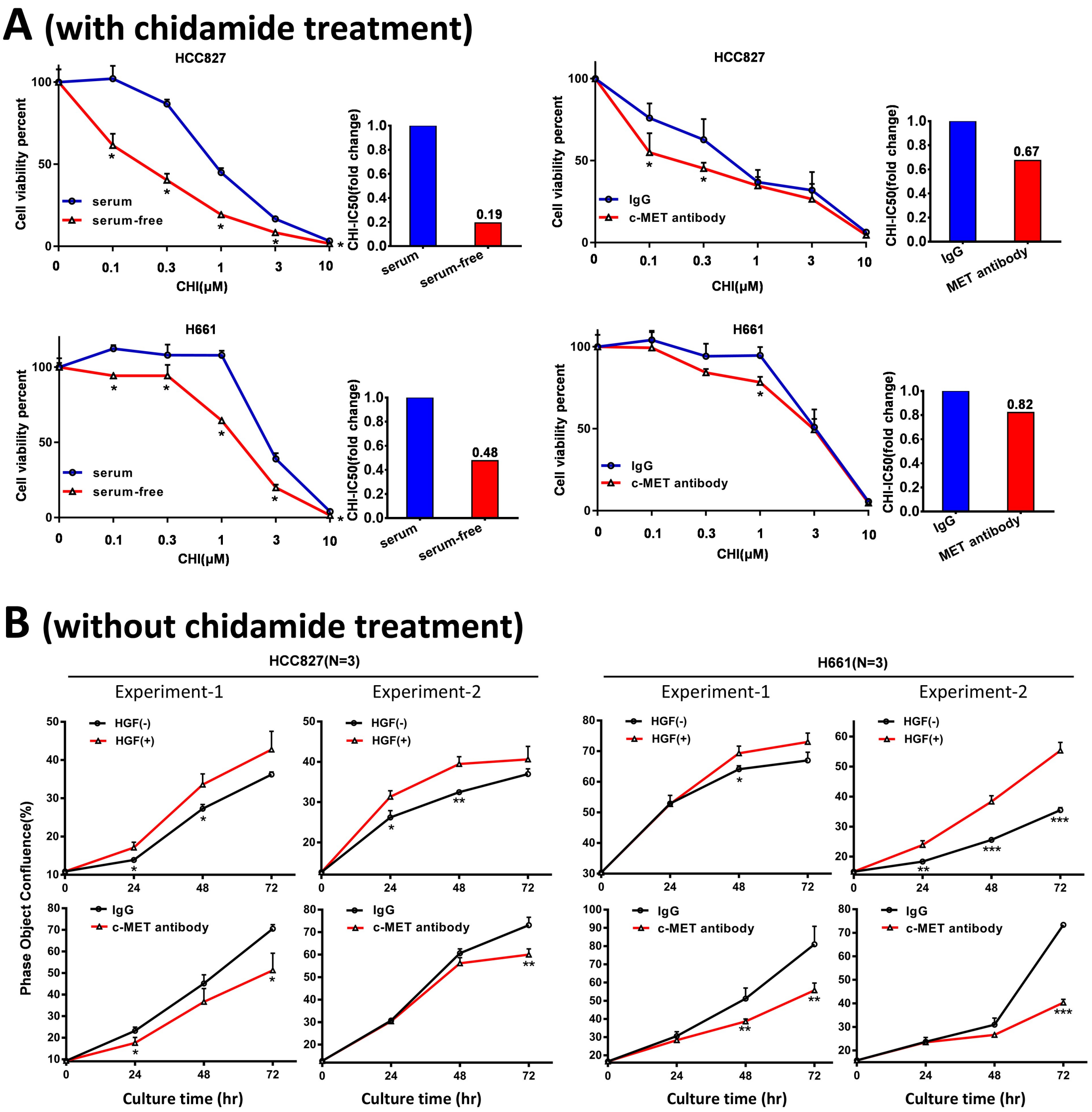


**Figure S2.** Effect of loss of function of *c-MET* by antibody (2.5 μg/mL) against c-MET or HGF-deprivation (serum-free) on the proliferation/viability of HCC827 and H661 cells treated with and without various concentrations of chidamide (CHI). The experimental conditions were the same as described in Figure 4 legend. (**A**) With chidamide treatment; The fold change of chidamide IC50 for cells treated with c-MET antibody or cultured in serum-free medium (HGF-deprivation) relative to cells treated with IgG control or cultured in serum-containing medium was labeled in right chart, respectively. (**B**) Without chidamide treatment. The results of two repeat experiments were illustrated. HGF(-),HGF-deprivation (serum-free); HGF(+), serum-free medium with HGF supplement

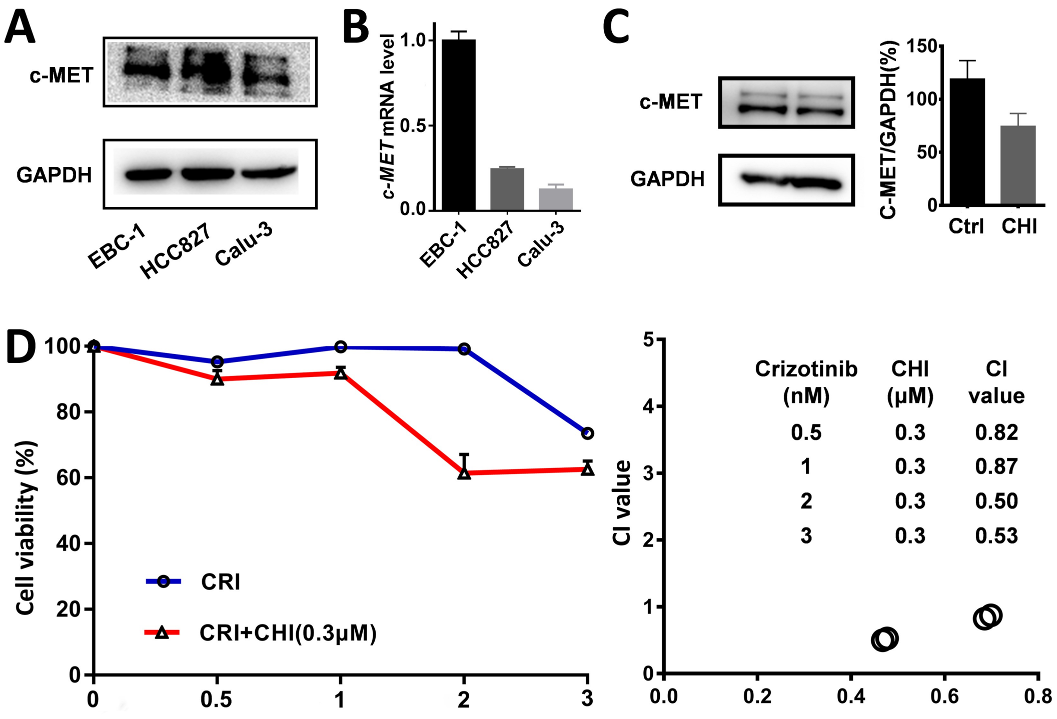


**Figure S3.** Synergistic effect of chidamide-crizotinib in NSCLC cell line EBC-1 with *c-MET* gene amplification. (**A** and **B**) The levels of baseline c-MET protein and mRNA in EBC-1 and other cell lines in Western blot and quantitative RT-PCR analyses, respectively. (**C**) The level of c-MET protein in EBC-1 cells with treatment of chidamide (CHI, at 1/4 IC50 [0.3μM]) for 72 hr in Western blotting. (**D**) Cell viability was measured by using the IncuCyte platform (left chart). The crizotinib-IC50 values for BEC-1 cells were calculated in the absence or presence of chidamide. The synergetic effect of chidamide-crizotinib cotreatment on cell proliferation inhibition was calculated using the CI equation and presented as Fa (fraction affected by the dose) in the Fa–CI plots (right chart);

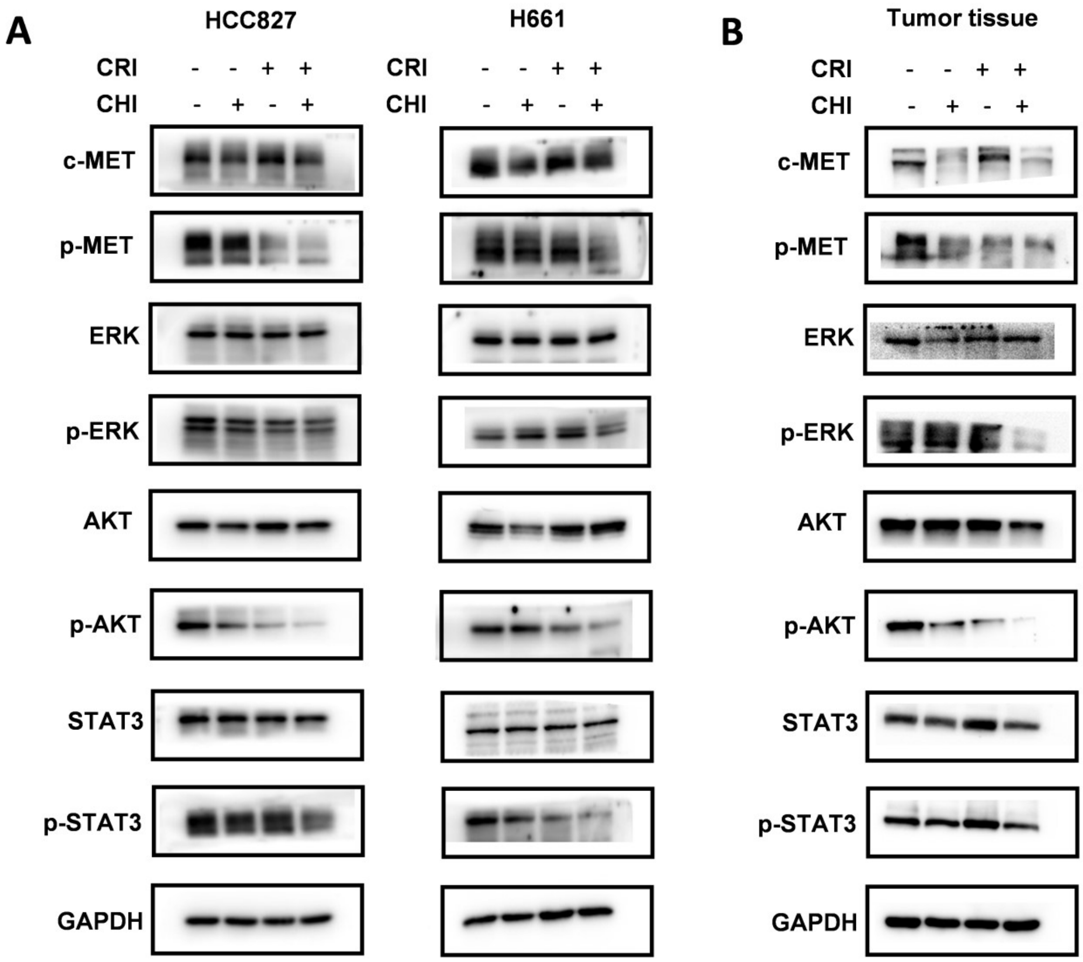


**Figure S4.** Effects of treatment with chidamide (CHI), crizotinib (CRI), or their combination on the phosphorylation levels of the receptor tyrosine kinase (RTK) signaling molecules MET, ERK, AKT and STAT3 proteins in NSCLC cells via Western blotting. (**A**) The NSCLC cell lines HCC827 and H661 treated with chidamide (0.25 μM for HCC827 and 1 μM for H661), crizotinib (1 μM), or their combination for 4 hr. (**B**) HCC827-derived tumors in mice treated with chidamide (5 mg/kg/d), crizotinib (25 mg/kg/d), or their combination for 21 days.

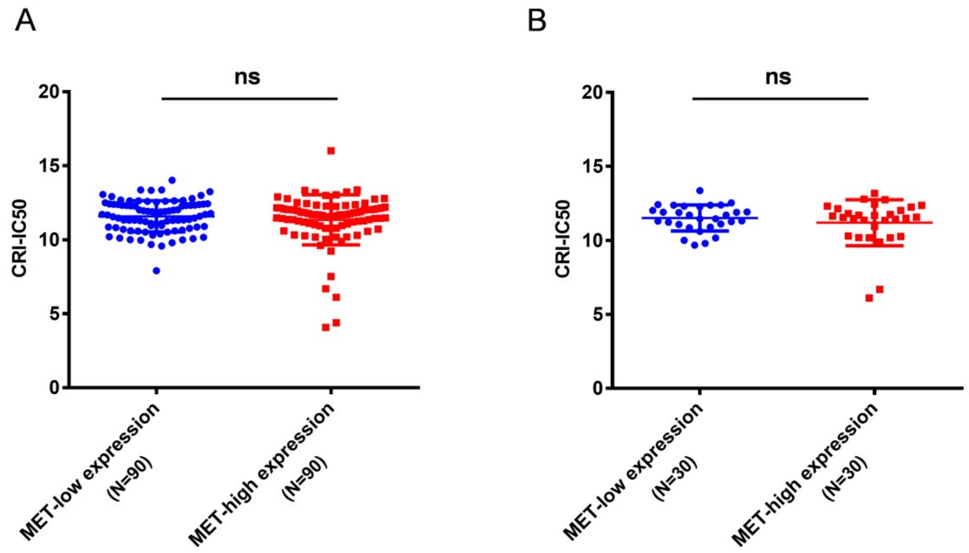


**Figure S5**. The IC50 values of crizotinib in human cancer cell lines with different expression statuses of *c-MET* mRNA in the Cancer Cell Line Encyclopedia (CCLE) project [26]. The cell lines were equally stratified into the high *c-MET* expression and low *c-MET* expression groups. (**A**) All 180 human cancer cell lines. (**B**) 60 lung cancer cell lines. ns, not significant.

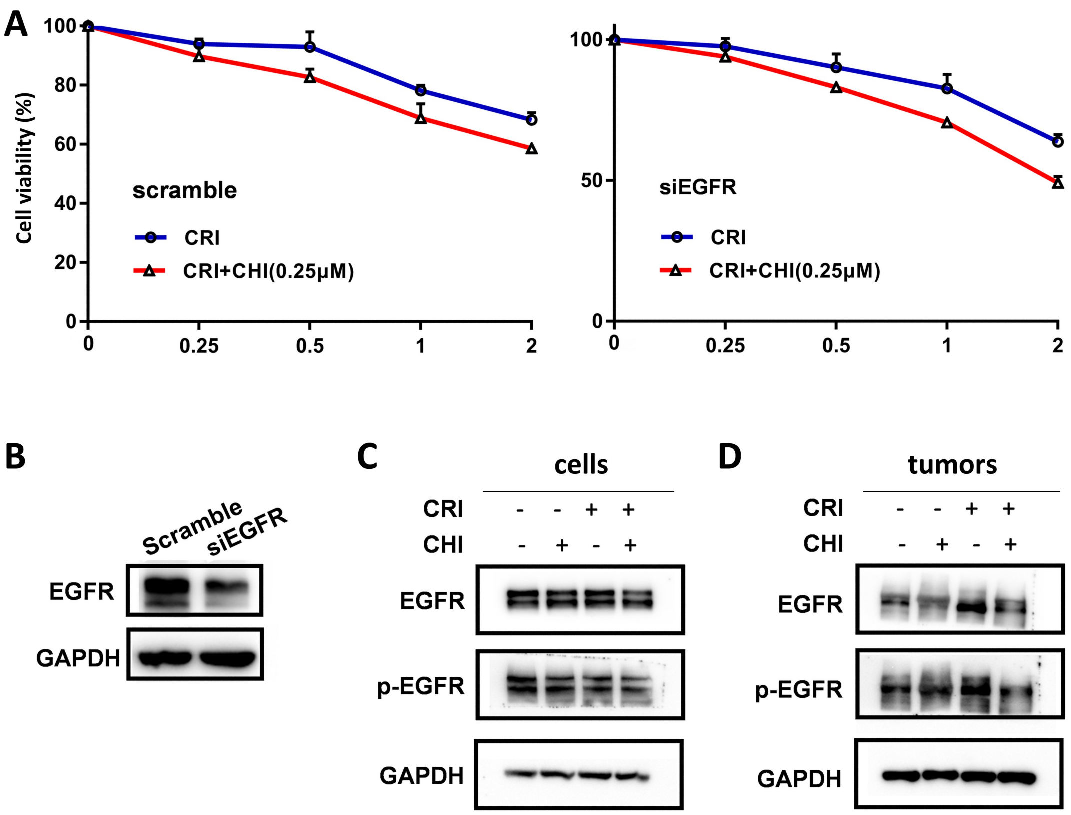


**Figure S6**. Effect of *EGFR* downregulation by siRNA on the viability and EGFR phosphorylation of HCC827 cells with treatment of chidamide, crizotinib, and their combination. (**A**) Synergistic effect of chidamide-crizotinib in HCC827 cells with and without siRNA knockdown of *EGFR* expression (siEGFR). The crizotinib-IC50 values for BEC-1 cells were calculated in the absence or presence of chidamide. (**B**) The level of global EGFR protein in HCC827 cells 48 hr posttransfection in Western blotting. (**C**) The levels of global EGFR and phosphorylated EGFR (pEGFR) proteins in HCC827 cells treated with chidamide (0.25 μM), crizotinib (1 μM), or their combination for 4 hr in Western blotting; (**D**) The level of global EGFR protein in HCC827-derived tumors in mice treated with chidamide (5 mg/kg/d), crizotinib (25 mg/kg/d), or their combination for 21 days.
